# Establishment of a healthy control iPSC line from an Eastern Indian donor as a population specific resource for disease modelling

**DOI:** 10.64898/2026.06.09.731103

**Authors:** Saileyee Roychowdhury, Vasanth Thamodaran, Deepika Joshi, Parimal Das

## Abstract

**Background:** iPSCs generated from healthy individuals constitute an important control resource for disease modelling applications but existing biobanks are highly skewed towards populations of European ancestry while well characterized control lines from Indian populations remain limited. Given the extensive genetic diversity of the Indian subcontinent, the availability of ethnically relevant healthy control lines is important for developing accurate disease models and reducing population specific confounding effects.

**Methodology:** We used peripheral blood mononuclear cells (PBMNCs) of a healthy female donor of Eastern Indian origin for the generation a wild type iPSC line using non-integrating episomal reprogramming vectors. Established colonies were expanded and characterized through morphological assessment, expression of pluripotency and trilineage markers, episomal vector clearance analysis, and chromosomal stability evaluation and mycoplasma contamination analysis.

**Results:** The line generated exhibited characteristic pluripotent stem cell morphology and also showed strong expression of pluripotency markers, was free from any contamination and free from the reprogramming vectors confirming an integration free system. The cells maintained a normal diploidy number during characterization. Expression of lineage specific markers associated with ectoderm, mesoderm and endoderm confirmed the developed iPSCs functional capacity to undergo trilineage differentiation.

**Conclusion:** We have developed and validated an iPSC line from an underrepresented Indian population. This well characterized, ethnicity specific iPSC line provides a valuable cell line for establishing a high quality, well characterized control baseline, which is a major missing element in South Asian stem cell repositories and thus will provide a solid foundation for future disease specific modelling and screening.

## 1. Introduction

The development of hiPSC technology has created new opportunities to study how disease develop, examine the underlying cellular mechanisms, and assess potential treatment strategies in human derived systems [1]. First described by Yamanaka et al. in 2006, hiPSCs are generated by reprogramming somatic cells through the expression of OCT4, SOX2, KLF4 and c-MYC or L-Myc [2], [3]. Like Embryonic stem cells, hiPSCs can proliferate indefinitely and can generate derivatives of ectodermal, mesodermal, and endodermal lineages, offering these advantages without many of the ethical concerns linked to embryo derived cells [4].

The impact of hiPSC technology has been particularly crucial in the field of neurodegenerative disease research [5]. Disorders like Amyotrophic lateral sclerosis (ALS), Spinal muscular atrophy (SMA), Parkinson’s disease (PD), and Alzheimer’s disease (AD) to name a few, remain difficult to study because access to living human neural tissue is extremely limited and traditional transgenic animal models often fail to fully reproduce human disease phenotypes. As a result, many findings from preclinical studies do not translate into clinical settings. hiPSC technology offers an alternative by enabling the generation of patient and donor specific cell types in vitro [6], [7]. Owing to their pluripotent nature, one of the key advantages of hiPSCs is their ability to generate a broad spectrum of human cell types, making them suitable for modelling disorders affecting multiple organ systems, including the neural, cardiac, hepatic and pancreatic lineages [8], [9]. This versatility has expanded their application beyond neurodegenerative disorders to a wide range of developmental, metabolic, cardiovascular and genetic diseases providing a more physiologically relevant system for studying various aspects of human diseases.

A major requirement for any hiPSC based disease model is the availability of well characterized healthy control lines. These controls provide the baseline against which disease associated phenotypes can be interpreted and help distinguish true disease related changes from normal genetic variation [10]. In addition to serving as reference standards, healthy control hiPSC lines are often used to establish differentiation protocols, optimize experimental conditions, and investigate fundamental aspects of human cellular biology. In situations, where patient derived hiPSC lines are unavailable or difficult to obtain, healthy control lines can also be used as a starting platform for disease modelling through targeted genetic engineering approaches. Enabling the introduction of specific disease associated variants into an otherwise genetically defined background [7].

However, most publicly available hiPSC resources originate from individuals of European and Caucasian ancestry, resulting in a significant lack of representation from other populations [11]. This is particularly relevant for the Indian subcontinent, which harbours extensive genetic diversity arising from thousands of ethnolinguistic and endogamous communities. Consequently, the Indian genome harbours unique genetic architectures, including distinct single nucleotide polymorphisms (SNPs), copy number variations, and a high burden of founder mutations that directly influence disease susceptibility, drug metabolism, and therapeutic efficacy [12]. As a result, the use of control lines derived from other ancestral background may not fully capture the genetic context of Indian populations and could overlook population specific modifiers or baseline cellular characteristics relevant to disease phenotypes [11].

Given that genetic background can affect disease manifestation and treatment responses, making ancestry matched controls therefore should be an important consideration for disease modelling studies. Despite the increasing use of hiPSCs in biomedical research, validated and well characterized iPSC resources from Indian donors to serve as a control platform remain exceptionally scarce[13]. Establishing a repository of ethnically relevant control lines is therefore essential for investigating complex disorders, including neurodegenerative diseases Furthermore, ensuring these lines are generated using integration free non viral methods, is vital for their long term genomic stability and downstream clinical application.

To address this gap, this study details the generation, characterization, and validation of a wild type iPSC line derived from the peripheral blood mononuclear cells (PBMNCs) of a healthy donor of Eastern Indian ancestry. By utilizing non integrating episomal reprogramming approach, we successfully generated a pluripotent, and genetically stable line which was subsequently and rigorously validated for its pluripotency potential, multi lineage differentiation capacity, and chromosomal integrity thereby establishing it as an ethnicity appropriate reference standard for Indian biomedical research.

## 2. Material and method

### 2.1. Healthy donor selection

A 30 year old healthy female donor of Eastern Indian origin with no reported personal or family history of neurological disorder was selected for the study. Written informed consent was obtained prior to sample collection, and the study was carried out in accordance with Institutional Ethics Committee approval. Before the extraction of peripheral blood, the donor underwent standard clinical screening to exclude virus-related infections, including HIV type I & II, as well as Hepatitis C to ensure biosafety and sample integrity for downstream applications.

### 2.2. PBMNC isolation and expansion of erythroid progenitor cells

About 5 ml of peripheral blood was collected in an EDTA coated vial for the isolation of peripheral blood mononuclear cells (PBMNCs). Isolation was performed using the Ficoll-Paque density gradient centrifugation method as described previously [14]. Briefly, the blood sample was diluted in a 1:1 ratio with CTS DPBS (without calcium and magnesium) (Gibco), and carefully layered onto a 5 ml of Ficoll-Paque premium (GE healthcare). Density gradient centrifugation was performed at 400 x g for 36 minutes at 18 °C to isolate PBMNCs. Following centrifugation, the PBMNC layer was collected and transferred to a fresh tube, followed by washing at 300 x g for 10 minutes. Residual erythrocytes were removed using RBC lysis buffer with incubation at 4 °C for 10 mins. The cells were subsequently pelleted by centrifugation and washed with 5 ml of CTS DPBS under the same centrifugation. The final PBMNC pellet, containing approximately 5 × 10^7^ cells, was resuspended in 1 ml of cryopreservation medium and stored at -80 °C for further use.

The following day, cryopreserved PBMNCs were thawed at 37 °C and immediately subjected to centrifugation at 300 x g for 10 mins to remove the residual cryoprotectant. Following centrifugation, the cell pellet was cultured in SFEM II (STEMCELL technologies) based erythroid expansion medium supplemented with SCF (50 ng/ml), IGF (40 µg/ml), EPO (3U/ml) and Il-3 (10 µg/ml). Cultures were maintained at 37 °C in a humidified 5% CO_2_ incubator for eight days to facilitate proliferation of erythroid progenitor cells.

### 2.3. Generation of iPSCs by episomal reprogramming

On day 9 of erythroid progenitor cell cultures derived from PBMNCs, reprogramming was initiated via integration free episomal plasmid nucleofection. A reprogramming plasmid cocktail was prepared comprising of electroporation buffer T and 1 μg each of pCXLE-hOCT3/4-shp53-f (Addgene, #27077), expressing *OCT3/4* and an shRNA against p53, 1ug of pCXLE-hSK (Addgene, #27078) expressing *SOX2* and *KLF4*, and 2 μg of pCXLE-hUL (Addgene, #27080) expressing L-Myc and LIN28. Erythroid progenitor cells were harvested, and approximately ∼1 x 10^6^ viable cells were counted and washed with calcium and magnesium-free DPBS. The cell pellet was resuspended in the plasmid-containing buffer and subjected to nucleofection using the Neon Nucleofection system under the following conditions: 1300 V, 30ms, 1 pulse. Post electroporation, the cells were immediately transferred to a 1% Geltrex-coated culture plate (Gibco, A33492-01) containing prewarmed erythroid expansion medium and maintained at 37°C with 5% CO_2_ to facilitate reprogramming and formation of iPSC colonies. The medium was changed on every alternate day from day 3 onwards with StemFlex medium (Gibco). On day 20, emerging colonies exhibiting characteristic features of hiPSCs were manually picked and expanded. For maintenance and further expansion, the cells were seeded in 1% Geltrex coated dishes containing prewarmed StemFlex medium and subsequently passaged using 1X EDTA (Invitrogen) solution. [14]

### 2.4. Determination of Pluripotency

For Pluripotency, iPSC colonies were dissociated using TrypLE™ Express (Thermo) solution and 40K single cell suspension was seeded in a 1% Geltrex coated plate with StemFlex medium. Once the cell reached around 40-50% confluency, they were taken for immunofluorescence studies.

### 2.5. Trilineage differentiation

To check the trilineage differentiation potential of the iPSCs the following protocols were used:

a. **Endoderm differentiation:** After dissociating the hiPSCs into single cell suspension using TrypLE™ Express solution, about 40K cells were seed onto a 1% Geltrex coated plates. Supplemented with StemFlex medium and Y-27632 (10µM) (Sigma, #SCMO75). in this medium, the cells were cultured or 48 hours post which StemFlex medium was replaced completely with endoderm differentiation medium containing: RPMI 1640 medium (Thermo,# 11875093), supplemented with FBS (0.2%), Activin A (25 ng/ml), CHIR 99021 (2 µM), NEAA (1×), Crotonate (5mM at pH 7.4) and cultured for 48 hours and subsequently taken for immunofluorescence studies[15].
b. **Mesoderm differentiation:** 40K cells from the single cell suspension were seeded onto a 1% Geltrex coated plate supplemented with StemFlex and Y-27632 (10µM). When the cells reached approximately 50% confluency, StemFlex medium was replaced completely with mesoderm differentiation medium consisting of Advanced RPMI, Glutamax (1×), CHIR 99021 (5 µM). the culture was maintained for 2 days and the medium was changed every alternate day, on day 3, cells were processed for immunofluorescence studies.
c. **Ectoderm differentiation:** 40K cells from single cell suspension was seeded onto a 1% Geltrex coated plate containing StemFlex medium and supplemented with Y-27632 (10µM). The next day, StemFlex medium was discarded completely and replaced with ectoderm differentiation medium containing Neurobasal media (Thermo, 21103049) and DF12 (Gibco) in 1:1 ratio along with Glutamax (1×), CHIR 99021 (3 µM), B27 (0.1%), Ascorbic acid (0.1mM), DMHI (2 µM), N2 (0.5%), and SB431542 (2 µM) and cultured for 6 days. Medium was changed every alternate day. On day 7, the cells were taken for immunofluorescence studies [16].

### 2.6. Immunocytochemistry

Cells were rinsed with 1X DPBS and subsequently fixed with 4% paraformaldehyde for 20 min at room temperature (RT), followed by washing the cells and permeabilization with 0.2% Triton X-100 in PBS for 15 min. After additional PBS washes, cells were incubated overnight at 4°C with the following primary antibodies specific for iPSC markers: rabbit anti-Nanog mAb, mouse anti-TRA-1-60 mAb, rabbit anti-OCT4 mAb, rabbit anti-SOX2 mAb, mouse anti-SSEA-4 mAb. mouse anti-TRA-1-81 mAb (1:500 dilution). To identify the trilineage potential of iPSC cells the following primary antibodies were used: rabbit anti-T-Brachyury mAb, rabbit anti-PAX6 mAb, rabbit anti-SOX17 mAb. (1:500) Following primary antibody incubation, cells were washed and incubated with following secondary antibodies: mouse anti-rabbit IgG Alexa Fluor 555, rabbit anti-mouse IgG Alexa Flour 488 (1:1000 dilution) for 2 hrs at RT on a rocker. Nuclei were counterstained with DAPI (1 µg/ml in PBS) for 15 mins. After a final PBS wash, stained cells were imaged using fluorescence microscopy (Olympus).

### 2.7. Karyotype analysis

GTG karyotyping was performed at passage 26 by Neuberg Anand Diagnostics, Bangalore to determine the chromosomal integrity and to exclude aneuploidy.

### 2.8. Mycoplasma testing

Mycoplasma contamination was assessed using MycoAlert™ assay kit (Lonza). Testing was performed on conditioned culture medium collected from the established line at passage 22, according to the manufacturer’s instructions. Briefly, the supernatant obtained from the cultured medium post centrifugation was mixed with MycoAlert reagent followed by incubation to get Read A. after another additional incubation with MycoAlert substrate, read B was obtained. Luminescence signal was quantified using a microplate reader (Tecan). Mycoplasma contamination was calculated from Read B/Read A ratio.

### 2.9. Episomal vector clearance

To confirm, episomal clearance, genomic DNA was extracted from the established line at passage 20 using DNA isolation Kit. PCR was performed with specific primers designed to amplify the vector backbone and plasmid DNA used for nucleofection served as the positive control, while a no template control was included as the negative control.

## 3. Results

### 3.1. Generation and morphological evaluation of the iPSC line

Following isolation from peripheral blood, PBMNCs were cultured in expansion medium to enrich the proliferative cell population prior to reprogramming. Initially the culture consisted of a heterogeneous mixture of round, non adherent cells (Fig 1) Over the following days, the cells progressively expanded and formed dense clusters of suspension cells (Fig. 2). Once sufficient cell numbers were obtained, the expanded PBMNCs were nucleofected with episomal reprogramming vectors. Post nucleofection of the donor PBMNCs, distinct cell colonies with well defined borders began to emerge (Fig. 3). After manual picking of colonies followed by their expansion in a feeder free culture, the cells exhibited characteristic morphological features of iPSCs, having well defined borders, a high nucleus: cytoplasm ratio (Fig 4). The line maintained with StemFlex medium on a 1% Geltrex coated plates with occasional dissociation with 1X EDTA solution, consistently expanded without spontaneous differentiation till culture 40.

**Fig 1.**
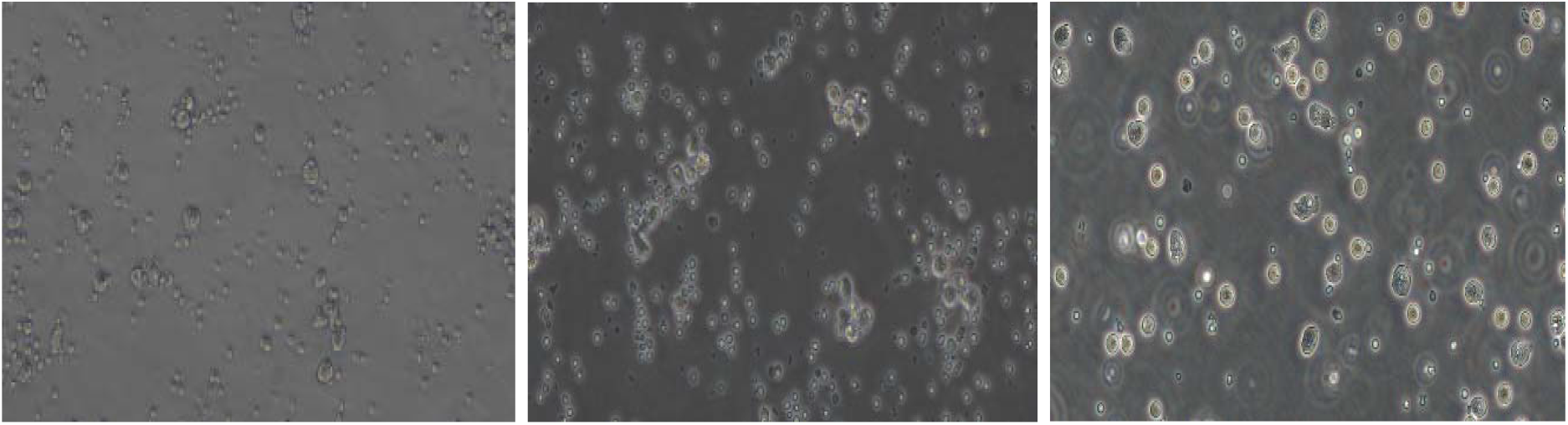
Morphological transition of donor PBMNCs in expansion culture at Days 5-Day 7 (20X magnification, Scale bar: 50 µM)

**Fig 2.**
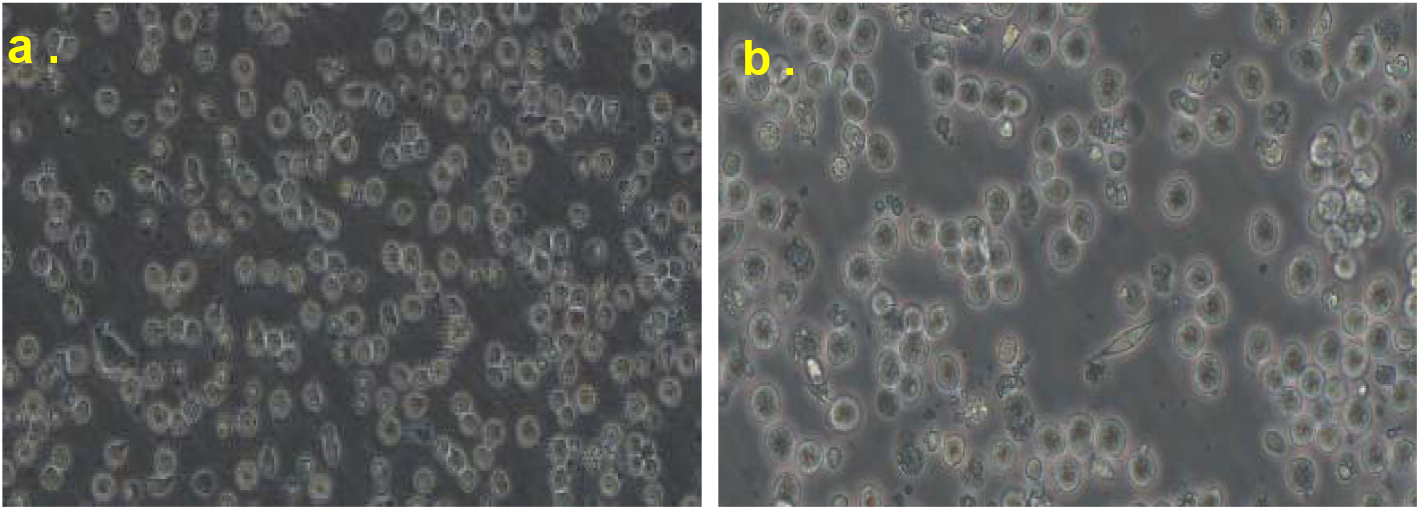
Progression of Erythroid progenitor cells post nucleofection with non integrating episomal vectors at (a) day 3 and (b) day 5 (20X magnification: Scale bar: 50 µM) capturing early cellular modelling

**Fig 3.**
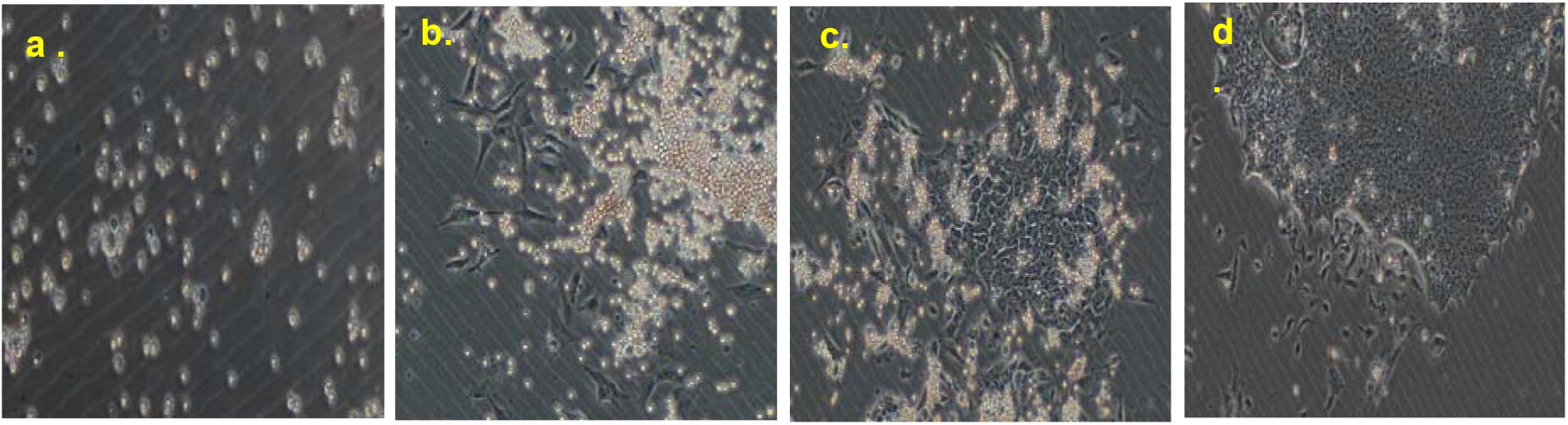
(a-d): Chronological timeline of the reprogramming process imaged at days 7,9,11,13 (10X magnification, scale bar: 100 µM), illustrating the gradual emergence of dense cell clusters transitioning into early packed cell aggregates.

**Fig 4.**
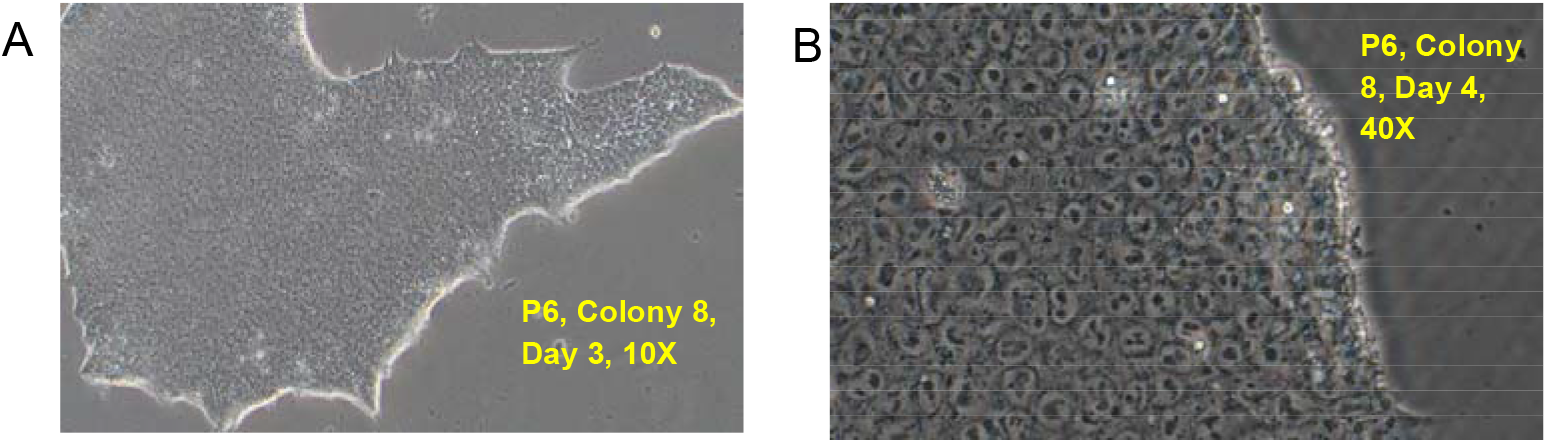
(A,B) Established hiPSC colonies at passage 6 under low and high power resolution (10X and 40X magnification, respectively). Colonies exhibit classic stem cell like morphology, characterized by sharp, prominent borders, tightly packed cells, high nuclear: cytoplasm ratios and distinct nucleoli.

### 3.2. Pluripotency validation by expression of markers

Immunofluorescence analysis of the developed hiPSCs confirmed strong expression of both core pluripotency markers (OCT4, SOX2, and NANOG) along with surface antigen markers (TRA-1-60, TRA-1-81, SSEA4); co-stained with nuclear maker DAPI, suggesting that the PBMNCs have successfully reprogrammed back into the pluripotent state

### 3.3. Confirmation of Trilineage differentiation potential

Immunofluorescence analysis of the differentiated cultures showed strong expression of SOX17 (specific to endoderm germinal layer), PAX6 (specific to Ectoderm germinal layer) and Brachyury (specific to mesoderm germinal layer), thereby suggesting the established hiPSC line has the full potential for trilineage differentiation (Fig 6).

### 3.4. Genomic stability analysis

To ensure the line is suitable for downstream modelling application we evaluated its genomic status. GTG banding karyotyping analysis showed normal, stable female diploid complement (46, XX) across all analyzed metaphase spreads, with no signs of gross chromosomal abnormalities including translocations, deletions or duplications induced by the reprogramming process (Fig 7).

### 3.5. Vector clearance analysis

To check the cells actually acquired pluripotent state and the expression of markers pluripotent markers are not due to the residual vector backbones used during reprogramming, PCR was performed targeting the episomal backbone. The amplified products when run on 1% agarose gel showed complete clearance of the vectors by passage 20 (Fig 8)

### 3.6. Mycoplasma testing

The line also tested negative for Mycoplasma contamination as suggested by the Read B/Read A ratio (Table 1).

**Table 1:** High sensitivity bioluminescent assay matrix validating the absolute absence of Mycoplasma contamination. Testing was done at passage 22.

| Sample | MycoAlert Reagent (A) | MycoAlert Substrate (B) | Result (B÷A) |
| --- | --- | --- | --- |
| Wild Type | 34 | 14 | 0.411 |

| Ratio | Interpretation of Results |
| --- | --- |
| <0.9 | Negative for mycoplasma |
| 0.9-1.2 | Quarantine cells and retest after 24 hrs |
| >1.2 | Mycoplasma contamination |

## 4. Discussion

The availability of well characterized healthy control iPSC lines is fundamental to the development of reliable stem cell based disease models [17]. While the use of hiPSCs have expanded rapidly over the past decade, control lines representing the Indian populations remain underrepresented in publicly available repositories [11], [18]. This gap becomes increasingly relevant when the genetic background can influence cellular phenotypes, susceptibility to disease associated molecular changes while studying a disease [11]. The generation of control lines from underrepresented populations therefore enhances the diversity of stem cell resources available for biomedical research.

In the present study, we successfully established a wild-type iPSC line from PBMNCs derived from the peripheral blood of a healthy donor of Eastern Indian origin (Fig.1-4). PBMNCs were selected as the somatic cell source due to their minimally invasive collection procedure and high reprogramming efficiency, which make them a preferable alternative to the more commonly used skin fibroblasts for iPSC cultures[19].

We chose to use an episomal expression system instead of lentiviral or retroviral delivery, which helps to minimize the well documented problems of random genomic integration of the transgene, including insertional mutagenesis, disruption of important endogenous loci, or unpredictable transgene reactivation during subsequent differentiation [20]. The confirmation of vector clearance (Fig. 8) suggested the phenotypic characteristics observed in this line can be considered as the natural genetic traits of the donor and not a trait resulting from artificial vector expression. This result is crucial if the generated line is intended to be used as a clean control.

The pluripotency of the line was well validated by robust nuclear expression of the pluripotency markers (Fig.5). Trilineage differentiation capacity was also demonstrated by the expression of relevant germ layer specific antibodies as shown in the immunofluorescence studies which will be useful in future differentiation application (Fig 6) and it was free from any Mycoplasma contamination (Fig.7).

**Fig 5.**
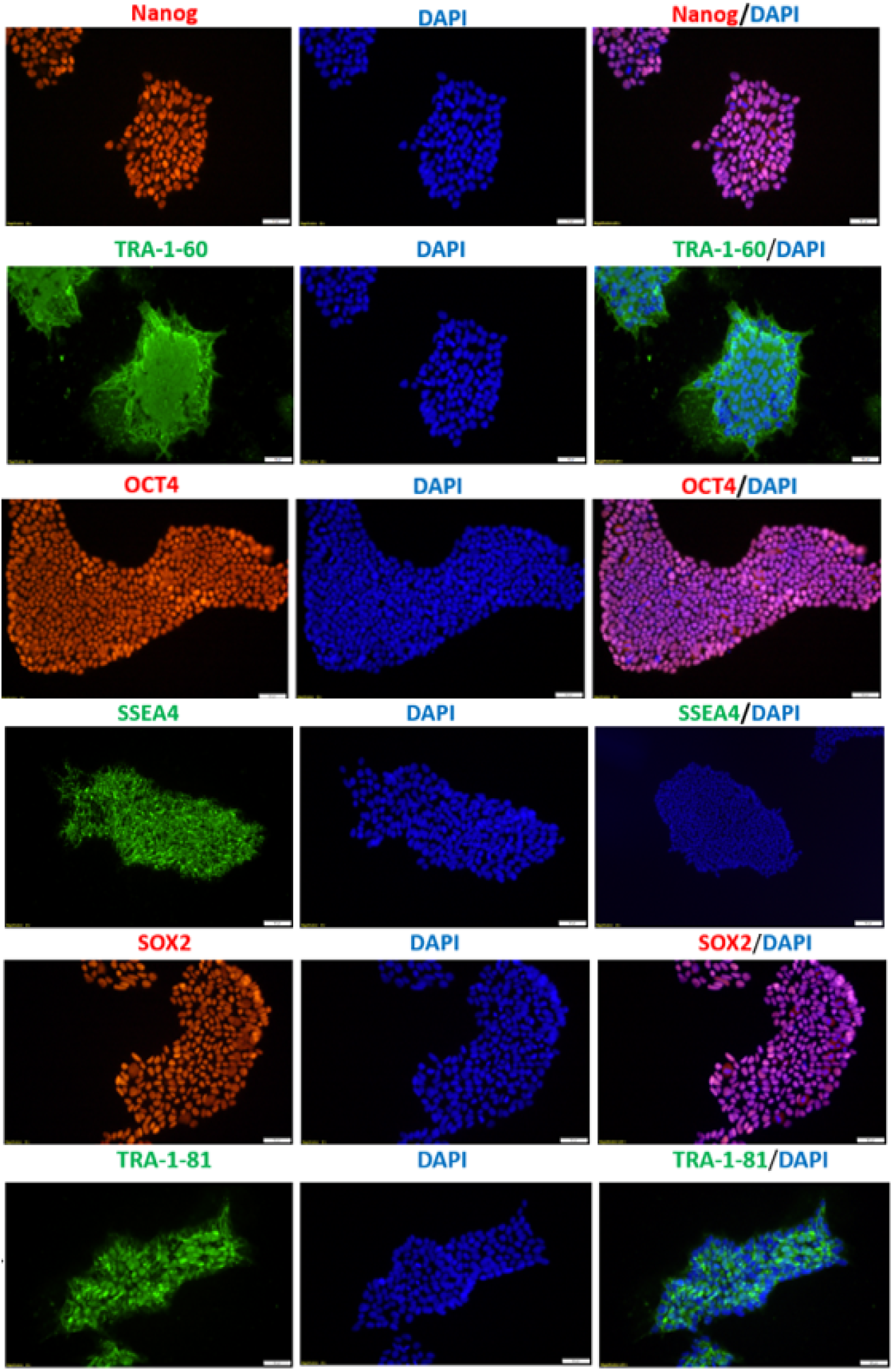
Immunofluorescence based pluripotency validation showing robust expression of core nuclear transcription factors (OCT4, SOX2, NANOG) and surface antigen markers (TRA-1-60, SSEA4, TRA-1-81); counterstained with DAPI to visualize nuclei. Scale bar: 50 µM

**Fig 6.**
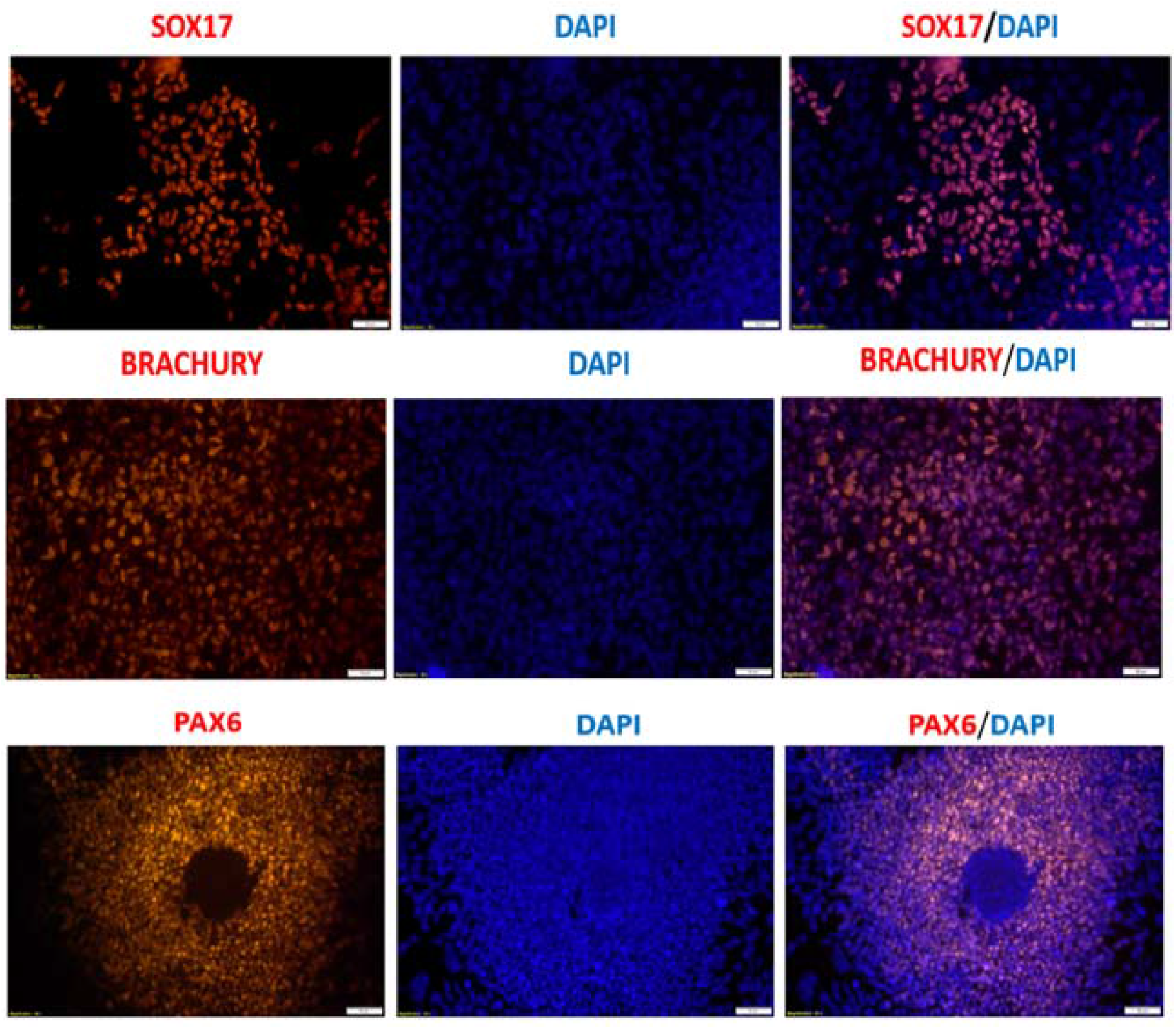
Immunofluorescence staining confirms differentiation potential of the iPSCs into all three embryonic germ layers: ectoderm (PAX6); mesoderm (Brachyury); and endoderm (SOX17). Scale bars: 50µM

**Fig 7.**
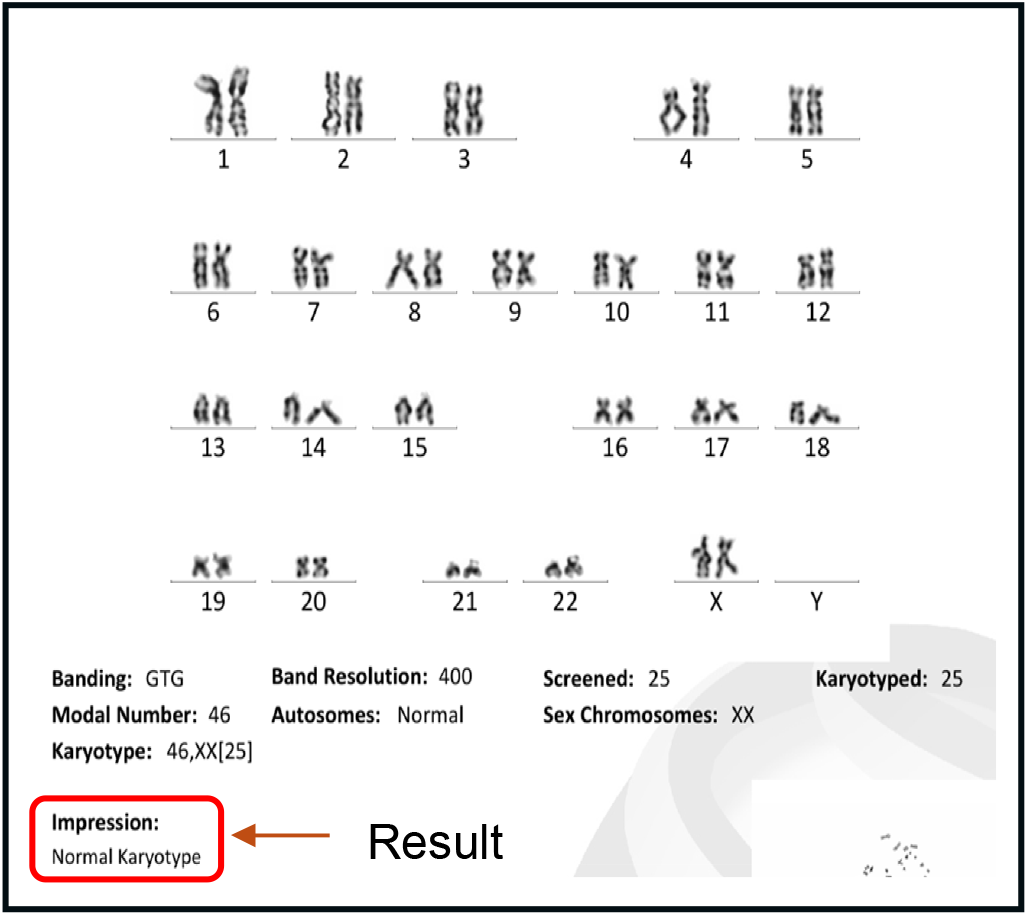
Representative GTG banding karyotypic profile of the established line at passage 26, demonstrating a normal, stable female diploid complement without detectable structural anomalies.

**Fig 8.**
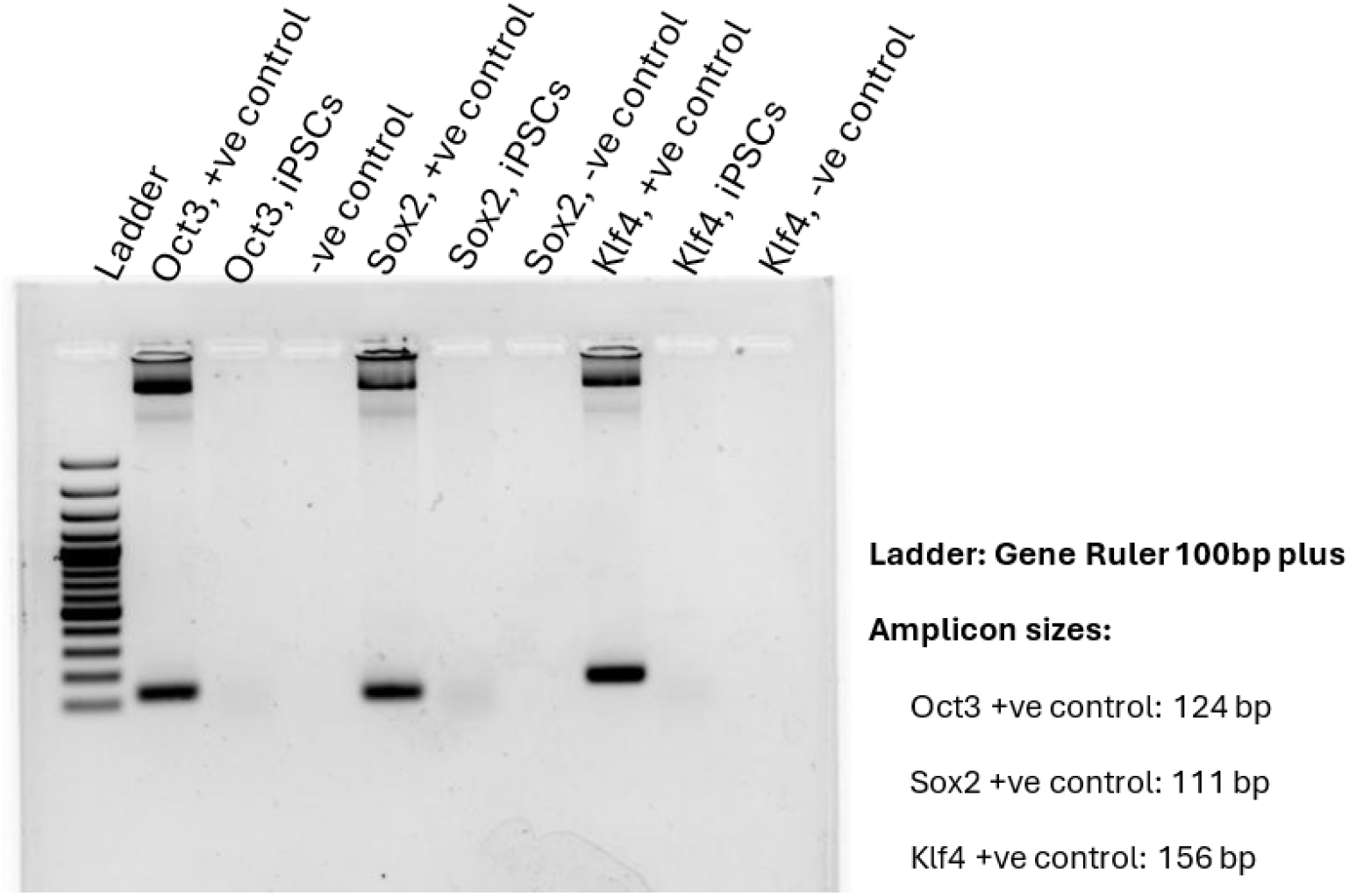
Genomic DNA isolated from established iPSC clones was subjected to PCR to assess the presence of residual episomal reprogramming vectors with Primer sets specific to episomal backbone elements were used. Plasmid DNA used for reprogramming served as a positive control for episomal presence, while a no-template control (NTC) was included as negative control. Absence of episomal-specific amplicons in the tested iPSC clones indicates successful clearance of reprogramming vectors, confirming their integration-free status. Representative agarose gel images show no detectable episomal bands in the analyzed clones at passage 20, whereas positive controls exhibit the expected amplicon sizes.

One of the most important considerations in the validation of any new control line is to establish its genomic stability. Chromosomal duplication or mosaicism and/or karyotypic anomalies can occur during long term cell culture and/or the stress of the reprogramming process itself[21]. Consequently, routine assessment of chromosomal stability remains an essential component of iPSC quality control. In our study, a normal 46, XX karyotype was preserved in the established cell line following multiple passages, therefore suggesting structural genomic integrity of this line (Fig. 7).

## 5. Conclusion

To conclude, this cell line is a physiologically relevant control with a genetically well defined background that better represents the Indian population, which is often underrepresented in the iPSC landscape globally. Population specific iPSC lines are essential for achieving a more comprehensive representation of the entire genetic diversity, thereby improving the accuracy and applicability of disease models and therapeutic strategies across diverse ethnic groups.

## Acknowledgement

The authors gratefully acknowledge the Stem Cell Facility and the laboratory members at the Tata Institute for Genetics and Society (TIGS), Bangalore, for providing the essential infrastructure, technical support, and scientific guidance that facilitated this research.

